# Reducing mitochondrial ribosomal gene expression does not alter metabolic health or lifespan in mice

**DOI:** 10.1101/2022.11.05.515295

**Authors:** Kim Reid, Eileen G. Daniels, Goutham Vasam, Rashmi Kamble, Georges E. Janssens, Man Hu, Alexander E. Green, Riekelt H. Houtkooper, Keir J. Menzies

**Author notes:** Correspondence – (RHH) and (KJM). co-first authors (Kim Reid, Eileen G. Daniels). co-last authors (Riekelt H. Houtkooper, Keir J. Menzies).

## Abstract

Maintaining mitochondrial function is critical to an improved health span and lifespan. Introducing mild stress by inhibiting mitochondrial translation invokes the mitochondrial unfolded protein response (UPR^mt^) and increases lifespan in several animal models. Notably, lower mitochondrial ribosomal protein (MRP) expression also correlates with increased lifespan in a reference population of mice. In this study, we tested whether partially reducing the expression of a critical MRP, *Mrpl54*, reduced mitochondrial DNA-encoded protein content, induced the UPR^mt^, and affected lifespan or metabolic health using germline heterozygous *Mrpl54* mice. Despite reduced *Mrpl54* expression in multiple organs and a reduction in mitochondrial-encoded protein expression in myoblasts, we identified few significant differences between male or female *Mrpl54*^+/-^ and wild type mice in initial body composition, respiratory parameters, energy intake and expenditure, or ambulatory motion. We also observed no differences in glucose or insulin tolerance, treadmill endurance, cold tolerance, heart rate, or blood pressure. There were no differences in median life expectancy or maximum lifespan. Overall, we demonstrate that genetic manipulation of *Mrpl54* expression reduces mitochondrial-encoded protein content but is not sufficient to improve healthspan in otherwise healthy and unstressed mice.

## INTRODUCTION

Since 1972, mitochondria have been implicated in the aging process^1-4^, initially as a source of free radicals in the mitochondrial theory of aging, and more recently as an indicator where reduced mitochondrial respiratory chain efficiency is considered a cellular hallmark of aging^5,6^. This is because mitochondrial dysfunction: (1) occurs during normal aging^7-9^; (2) is implicated in age-related pathologies such as type 2 diabetes, cardiovascular disease, and cancer^10-12^; and (3) aggravation of mitochondrial dysfunction accelerates aging^13^. If mitochondrial dysfunction contributes to aging, then improving mitochondrial function should slow normal aging^14^. Although seemingly contradictory, mitochondrial stress early in development maintains mitochondrial function with age in several animal models^15-19^. This stress triggers a compensatory beneficial response – i.e., a mitochondrial hormetic (mitohormetic) response, which is suggested to improve healthspan or lifespan^6,20^. If mild mitochondrial stress improves healthspan or extends lifespan in mammals, then this opens new possibilities in human healthspan and longevity research.

Specifically, mitochondrial translational stress is emerging as a promising mitohormetic strategy^15,21,22^. The mitochondrial ribosome (mitoribosome), which consists of ∼80 mitoribosomal proteins (MRPs)^23^, conducts mitochondrial translation to produce essential electron transport chain (ETC) complex protein subunits encoded by mitochondrial DNA. Previously, we showed that partial knockdown of any individual MRP, and the subsequent disruption of mitochondrial translation, improved metabolic health and extended the lifespan of *Caenorhabditis elegans*^15^. Reducing the expression of a single MRP induced a mitonuclear protein imbalance, a stoichiometric imbalance between nuclear-(nDNA) and mitochondrial-DNA (mtDNA) encoded ETC subunits. This imbalance subsequently activated the mitochondrial unfolded protein response (UPR^mt^) – a nuclear transcriptional response that maintains or restores mitochondrial proteostasis^24-26^. Strikingly, the variation in expression of MRP genes amongst recombinant-inbred BXD mouse strains^27,28^, e.g., *Mrps5* expression, inversely correlated with a ∼250-day increase in lifespan^15^. Further, fibroblast cell cultures from long-lived Snell dwarf mice had greater expression of UPR^mt^ markers, including elevated 60kDa mitochondrial heat shock protein (HSP60 or HSPD1)^29^. Therefore, chronic disruption of mitochondrial translation and induction of the UPR^mt^ may promote both longevity and an improved healthspan via mitohormesis^15,30-35^.

The observation that the reduction of MRP expression and consequent induction of mitonuclear imbalance induced the UPR^mt^ and promoted longevity across species led us to a general hypothesis: partial disruption of mitochondrial translation in a mouse model would result in improved metabolic health with age and/or an increased lifespan. To test this hypothesis, we engineered a whole-body mouse model heterozygous for the *Mrpl54* gene (*Mrpl54*^*+/-*^). The MRPL54 protein was selected because: (1) reduced expression of *Mrpl54* invokes the UPR^mt^ and extends *C. elegans* lifespan^15^; (2) MRPL54 is evolutionarily conserved between species^36^; and (3) MRPL54 is essential to both recruitment of essential factors for mitochondrial translation^37,38^ and ETC function^39^. To assess metabolic health with age, we subjected both male and female *Mrpl54*^+/-^ and *Mrpl54*^*+/+*^ (wildtype, WT) littermate mice to *in vivo* metabolic phenotyping at 6-, 18-, and 24-months of age. In parallel, we conducted a longevity study to assess the effect of mitochondrial stress on life expectancy and lifespan.

## RESULTS

### Generation of Mrpl54^+/-^ mice

To investigate the effect of reduced *Mrpl54* expression on whole body metabolism and longevity, we generated mice heterozygous for *Mrpl54* (Fig. 1a) by deleting exon 2 in the *Mrpl54* gene. Homozygous *Mrpl54* exon 2 deletion proved lethal, as no *Mrpl54*^*-/-*^ mice were observed at weaning, yet WT and *Mrpl54*^*+/-*^ were born at the expected Mendelian ratio (Fig. 1b). Next, to assess if *Mrpl54*^*+/-*^ led to reduced expression of the targeted gene, we performed qPCR analysis in gastrocnemius, heart, liver, interscapular brown adipose tissue (BAT), and proximal colon tissues from 7-week-old male and female mice. We found that the relative expression of *Mrpl54* was highest in gastrocnemius and lowest in the liver (Fig. 1c). Additionally, in all tissues tested, *Mrpl54* expression was reduced by ∼50% in *Mrpl54*^+/-^ mice compared to WT (Fig. 1d).

**Figure 1.**
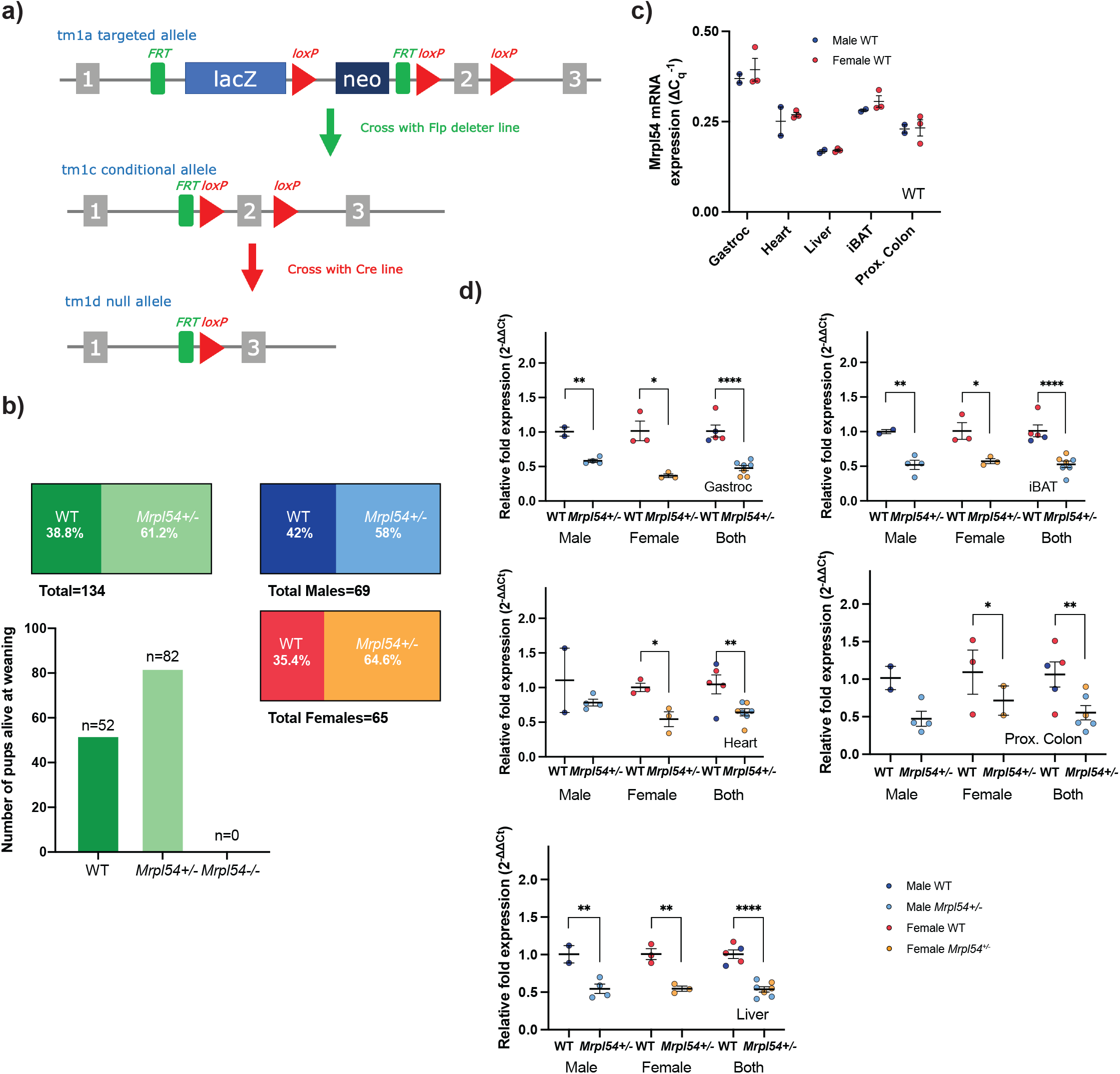
Generation and basic characterization of the *Mrpl54*^*+/-*^ model. **a)** Generation of the *Mrpl54*^*+/-*^ mouse model through the deletion of exon 2 at the *Mrpl54* allele. Adapted from the Mouse Genome Database^58^ **b)** The number of WT and *Mrpl54*^*+/-*^ pups born to 28 *Mrpl54*^*+/-*^ breeding pairs followed a Mendelian ratio for homozygous lethality. **c)** Relative *Mrpl54* gene expression in tissues from 7-week-old WT males (blue, n=2) and females (orange, n=3) as determined by RT-qPCR. Values were normalized to 60S acidic ribosomal protein P0 (*36b4* or *RPLP0*) and beta-2-microglobulin (*B2m*) expression. **d)** Reduced *Mrpl54* gene expression (∼50%) in tissues derived from 7-week-old *Mrpl54*^*+/-*^ males (light blue, n=2) and females (light orange, n=3) compared to WT male (blue, n=4) and female (orange, n=3). Values were normalized to *36b4* and *B2m* expression. Graphs show mean ± SEM, *P ≤ 0.05, **P ≤ 0.01, and ****P ≤ 0.0001

### No major metabolic phenotype in adult Mrpl54^+/-^ mice

To examine the effect of reduced *Mrpl54* expression on metabolic health in adult mice, we subjected male and female *Mrpl54*^+/-^ and WT mice to a comprehensive metabolic phenotyping protocol (Fig. S1). In both males and females in the combined longevity and metabolic study cohorts (the latter study animals included up until entering metabolic phenotyping, as described in methods), there were no differences in body mass between *Mrpl54*^+/-^ and WT mice (Fig. 2a). Both genotypes had similar lean- or fat-mass body composition across genotypes in males; although, females in the metabolic health study at 6 months (6M) had slightly lower lean and fat mass (Fig. 2b). Metabolic indices, examined via indirect calorimetry, exhibited no differences in mean O_2_ volume (VO_2_) (Fig. 2c) or mean respiratory exchange ratio (RER) (Fig. 2d). Mean VO_2_ was lower in 6M female *Mrpl54*^+/-^ animals during the less active light phase; however, an ANCOVA analysis^40^ attributed this difference to their reduced body mass (Fig 2c inset). Male spontaneous ambulatory activity in testing cages was not different, yet there was an increase in activity in 6M *Mrpl54*^+/-^ females when compared to WT (Fig. 2e), possibly in line again with the reduced body weight and VO_2_. Like the increase in ambulatory activity of female *Mrpl54*^+/-^ mice, 6M females exhibited improved running capacity in a forced exercise treadmill test (Fig. 2f). Whole body glucose handling was not different between genotypes during an oral glucose tolerance test (OGTT; Fig. 2g) or an intraperitoneal insulin tolerance test (iITT; Fig. 2h) in 6M male and female mice. Since skeletal muscle exhibited the highest relative levels of *Mrpl54* gene expression and BAT tissue the second highest (Fig. 1c), and because of their importance for non-shivering and shivering thermogenesis, we evaluated potential differences in metabolic heat production by exposing mice to an acute cold challenge (4°C). Mice from both genotypes maintained their core body temperature similarly for four hours for 6M males and females (Fig. 2i). At sacrifice, the masses of the heart, spleen and kidney were reduced as a percentage of body mass in *Mrpl54*^+/-^ males with no changes in skeletal muscle, brain, or liver (Fig. 2j). In females, there were no changes in organ masses expressed as a percentage of body mass. Overall, there were no robust morphological or metabolic changes in adult *Mrpl54*^+/-^ mice apart from the increased physical activity in female *Mrpl54*^+/-^ mice.

**Figure 2.**
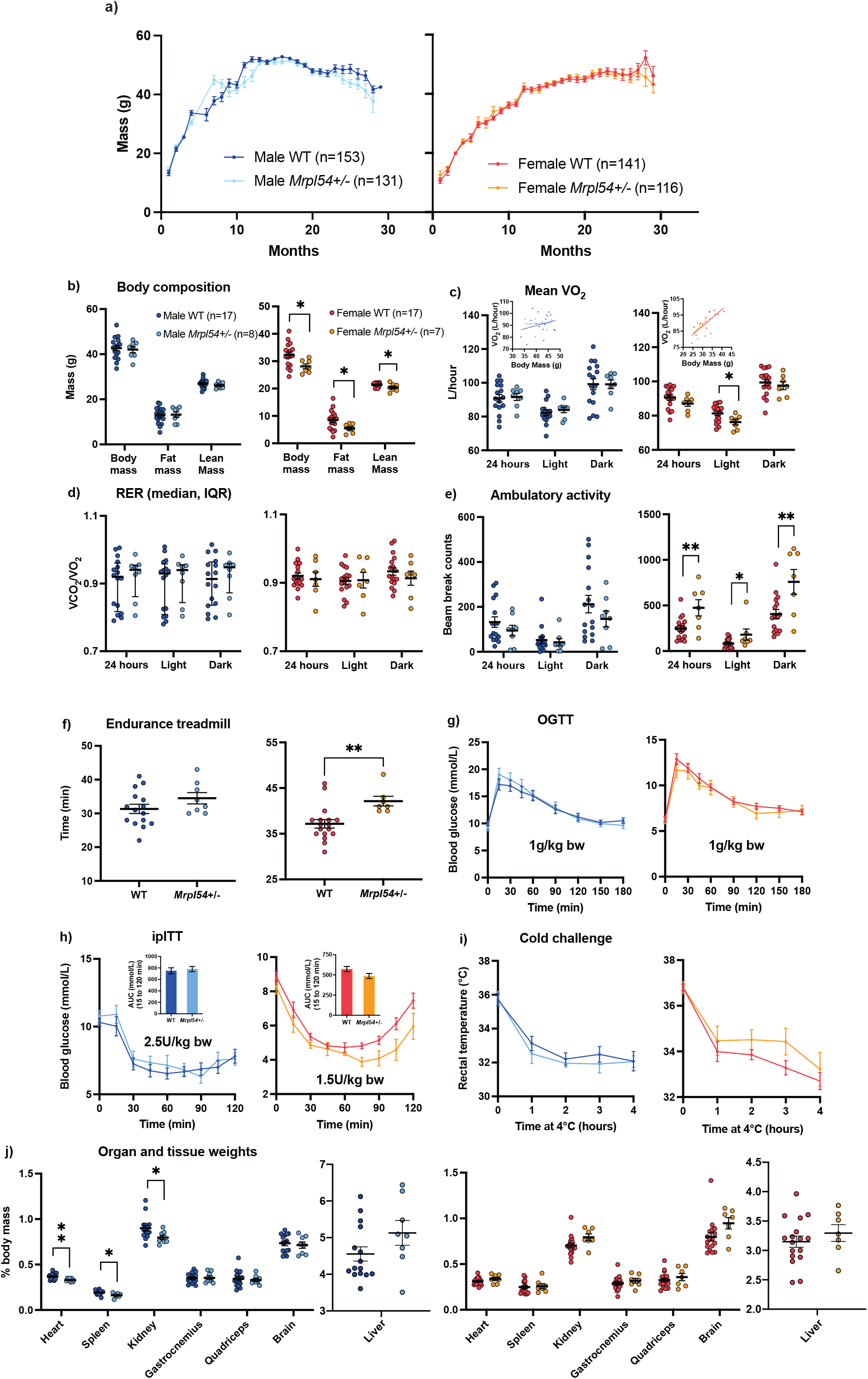
Metabolic phenotyping in 6-month-old male and female *Mrpl54*^*+/-*^ and wild type mice. **a)** No differences in body mass (g) between either male *Mrpl54*^*+/-*^ (n=131) and WT (n=153) or female *Mrpl54*^*+/-*^ (n=116) and WT (n=141) over the course of total lifespan. **b)** Body composition (g) by EchoMRI in 6M adult mice. The left graph shows no difference in total body, fat, or lean mass between *Mrpl54*^*+/-*^ and WT males. The right graph shows reduced body, fat, and lean mass in *Mrpl54*^*+/-*^ females compared to WT. **c)** Mean VO_2_ (L/hour) by CLAMS in 6M adult mice. Left panel shows no differences in VO_2_ for 24-hour, light, or dark periods between *Mrpl54*^*+/-*^ and WT males. Right panel shows a lower VO_2_ during the light cycle for *Mrpl54*^*+/-*^ females compared to WT, with no difference for the 24-hour or dark period. ANCOVA results (insets) indicate the difference in mean female VO_2_ are explained by the difference in female body mass. **d)** Mean RER by CLAMS in 6M adult mice showed no differences between *Mrpl54*^*+/-*^ and WT in either males or females. **e)** Mean ambulatory motion (beam break counts) in 6M adult mice. Left graph shows no differences in ambulatory motion for 24-hour, light, or dark periods between *Mrpl54*^*+/-*^ and WT males. Right graph shows an increase in ambulatory motion for the 24-hour, light, and dark periods in *Mrpl54*^*+/-*^ females compared to WT. **f)** Mean duration (minutes) on an endurance treadmill for 6M adult mice. The left graph shows a trend towards an increase in endurance time for *Mrpl54*^*+/-*^ males (n=8) compared to WT (n=15). The right graph shows an increase in endurance time for *Mrpl54+/-* females compared to WT. **g)** Mean blood glucose levels (mmol/L) over time in response to an oral glucose challenge (OGTT) in 6M adult mice showed no differences in blood glucose levels between *Mrpl54*^*+/-*^ and WT in either males or females. **h)** Mean blood glucose levels (mmol/L) over time in response to intraperitoneal insulin challenge (iITT) in 6M adult mice showed no differences in blood glucose levels or AUC (insets) between *Mrpl54*^*+/-*^ and WT in either males or females. **i)** Mean rectal temperature (°C) in response to a 4-hour 4°C cold challenge in 6M adult mice showed no differences in rectal temperature over time in males (WT, n=16 vs. *Mrpl54*^*+/-*^, n=7) or females. **j)** Relative organ weights (% body mass) in 6M adult necropsied mice. The left graph shows a decrease in the relative mass of the heart, spleen, and kidney in *Mrpl54*^*+/-*^ males (n=7-8) compared to WT (n=14-15). The right graph shows no differences in the relative masses of organs in *Mrpl54*^*+/-*^ females compared to WT. Graphs show mean ± SEM, *P ≤ 0.05, **P ≤ 0.01, and ****P ≤ 0.0001. Unless otherwise indicated, the number analysed are as follows: male WT (blue, n=16); male *Mrpl54*^*+/-*^ (light blue, n=8); female WT (orange, n=17); and female *Mrpl54*^*+/-*^ (light orange, n=7).

### Aged Mrpl54^+/-^ mice did not show improved metabolic phenotypes

Given that disruption of mitochondrial translation and/or UPR^mt^ induction has been linked to improved healthspan in several animal models^15,30-35^, we examined whether reduced *Mrpl54* expression altered whole body metabolism during the aging trajectory. Female and male *Mrpl54*^+/-^ and WT mice were aged 24 months (24M) before performing the same metabolic phenotyping tests used for adult 6M mice. While there were no differences in body composition between *Mrpl54*^*+/-*^ and WT, female mice had increased fat mass and decreased lean mass with age, as a proportion of body mass, while the opposite was observed in males – fat mass decreased and lean mass increased with age, as a proportion of body mass (Fig. S2, Fig. 3a). Like 6M animals, we did not observe any differences in indirect calorimetry measurements between *Mrpl54*^*+/-*^ and WT at 24M (Fig. 3b-d). At 24M, there were no differences in running capacity (Fig. 3e), which contrasts our observations in 6M *Mrpl54*^*+/-*^ female mice where running capacity was higher (Fig. 2e) but can be attributed to the difference in body weight at 6M. In the 24M mice, the response to OGTT was similar between *Mrpl54*^*+/-*^ and WT (Fig. 3f), as was the response to the iITT (Fig. 3g). There were no differences between genotypes when exposed to a cold challenge at 24M (Fig. 3h). There were no differences in heart rate, diastolic blood pressure, or systolic blood pressure between genotypes for either males or females at 24M (Fig. 3i). Finally, at 24M there were no differences in organ mass as a percentage of body mass (Fig. 3j). The 18M cohort displayed the same results as the 24M cohort for both males and females (Fig. S3a-h, S3j); except 18M male *Mrpl54*^+/-^ mice demonstrated a mean diastolic blood pressure that was higher than WT (Fig. S3i). Altogether, aging *Mrpl54*^+/-^ mice did not reveal robust differences from WT in metabolic parameters.

**Figure 3.**
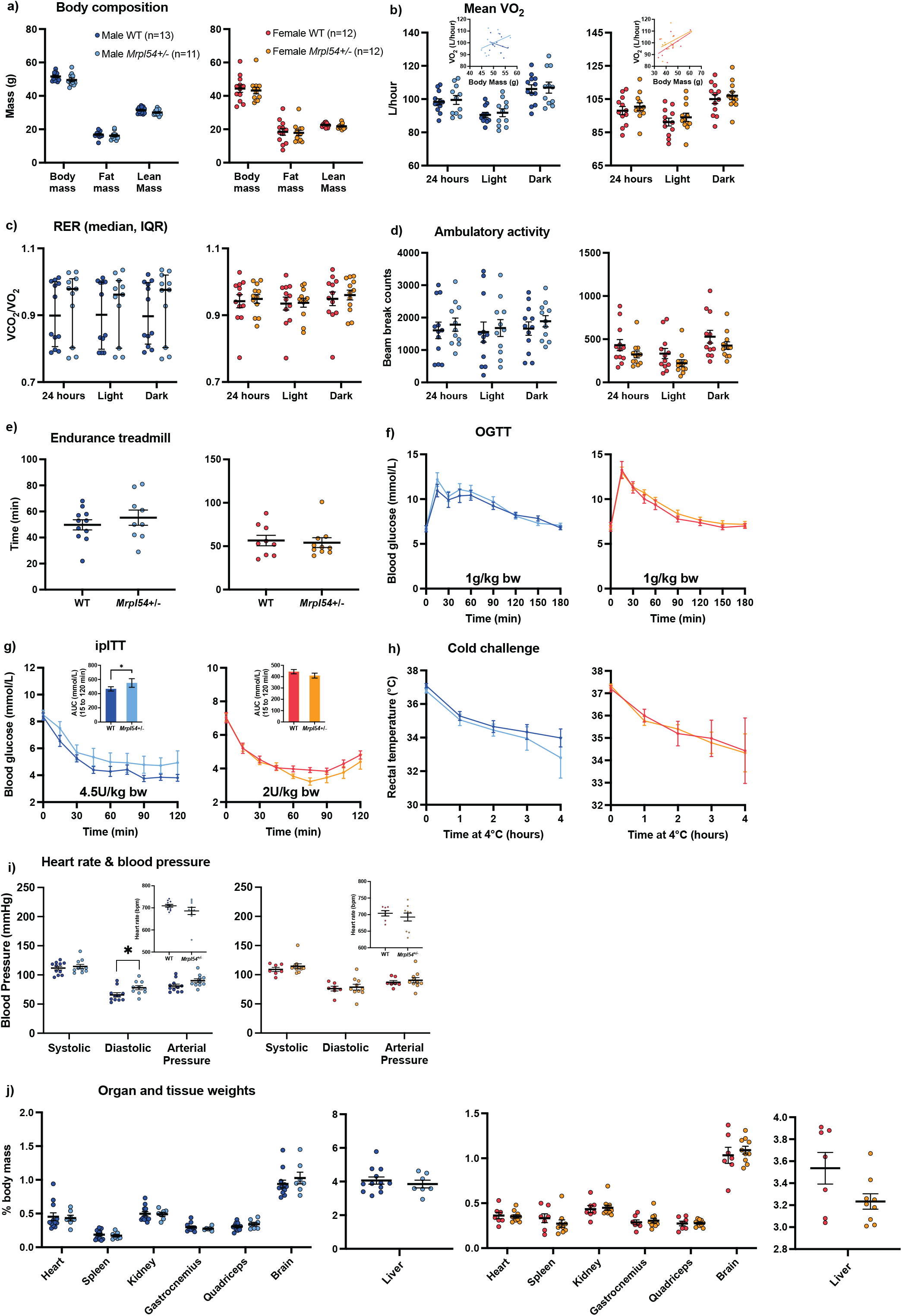
Metabolic phenotyping in 24M *Mrpl54*^*+/-*^ and wild type mice. **a)** Body composition (g) by EchoMRI in 24M old mice showed no difference in total body, fat, or lean mass between either male *Mrpl54*^*+/-*^ (n=12) and WT (n=10) or female *Mrpl54*^*+/-*^ (n=11) and WT (n=13). **b)** Mean VO_2_ (L/hour) by CLAMS and body mass ANCOVA analysis (insets) in 24M old mice showed no differences for 24-hour, light, or dark periods between either male *Mrpl54*^*+/-*^ (n=10) and WT (n=9) or female *Mrpl54*^*+/-*^ (n=11) and WT (n=13). **c)** Mean RER by CLAMS in 24M old mice showed no difference between either male *Mrpl54*^*+/-*^ (n=10) and WT (n=9) or female *Mrpl54*^*+/-*^ (n=11) and WT (n=13). **d)** Mean ambulatory motion (beam break counts) in 24M old mice showed no difference between either male *Mrpl54*^*+/-*^ (n=9) and WT (n=9) or female *Mrpl54*^*+/-*^ (n=10) and WT (n=13). **e)** Mean duration (minutes) on an endurance treadmill for 24M old mice showed no difference in endurance time between either male *Mrpl54*^*+/-*^ (n=6) and WT (n=7) or female *Mrpl54*^*+/-*^ (n=7) and WT (n=9). **f)** Mean blood glucose level (mmol/L) over time in response to an oral glucose challenge (OGTT) in 24M old mice showed no difference between either male *Mrpl54*^*+/-*^ (n=10) and WT (n=8) or female *Mrpl54*^*+/-*^ (n=10) and WT (n=12). **g)** Mean blood glucose level (mmol/L) and AUC (insets) over time in response to intraperitoneal insulin challenge (iITT) in 24M old mice showed no differences between either male *Mrpl54*^*+/-*^ (n=8) and WT (n=8) or female *Mrpl54*^*+/-*^ (n=10) and WT (n=11). **h)** Mean rectal temperature (°C) in response to a 4-hour 4°C cold challenge in 24M old mice showed no difference between either male *Mrpl54*^*+/-*^ (n=4) and WT (n=3) or female *Mrpl54*^*+/-*^ (n=4) and WT (n=7). **i)** Mean heart rate (beats/min) and blood pressure (mmHg) in 24M old mice showed no difference between either male *Mrpl54*^*+/-*^ (n=6) and WT (n=5) or female *Mrpl54*^*+/-*^ (n=5) and WT (n=8); although, there is a trend for higher diastolic pressure in male *Mrpl54*^*+/-*^ mice compared to WT. **j)** Relative organ weights (% body mass) in 24M necropsied mice showed no differences between either male *Mrpl54*^*+/-*^ (n=4-5) and WT (n=3-4) or female *Mrpl54*^*+/-*^ (n=3) and WT (n=6-7). Graphs show mean ± SEM; male WT (blue); male *Mrpl54*^*+/-*^ (light blue); female WT (orange); and female *Mrpl54*^*+/-*^ (light orange).

### Lifespan and life expectancies were not altered in Mrpl54^+/-^ mice

Since reduced MRP gene expression extended lifespan in *C. elegans* and correlated with long lifespan in BXD mouse strains, we tested whether reducing *Mrpl54* expression affected lifespan or life expectancy in mice. We followed *Mrpl54*^*+/-*^ male and female mice until they reached a humane endpoint or a natural death and assessed mean and 24-month life expectancy and maximum lifespan. We included 24-month life expectancy because strain-specific conditions and diseases affect C57BL/6 mice after the age of 24-months, which can confound results of aging studies^41^. We observed no significant differences in median life expectancy, 24-month life expectancy, or maximum lifespan between *Mrpl54*^+/-^ and WT, for either males or females (Fig. 4a); although, median and 24-month life expectancy for *Mrpl54*^+/-^ males trended higher compared to WT (median life-expectancy 705 days vs. 679 days: 24-month hazard ratio [HR] 1.38, 95% CI 0.93, 2.03).

**Figure 4.**
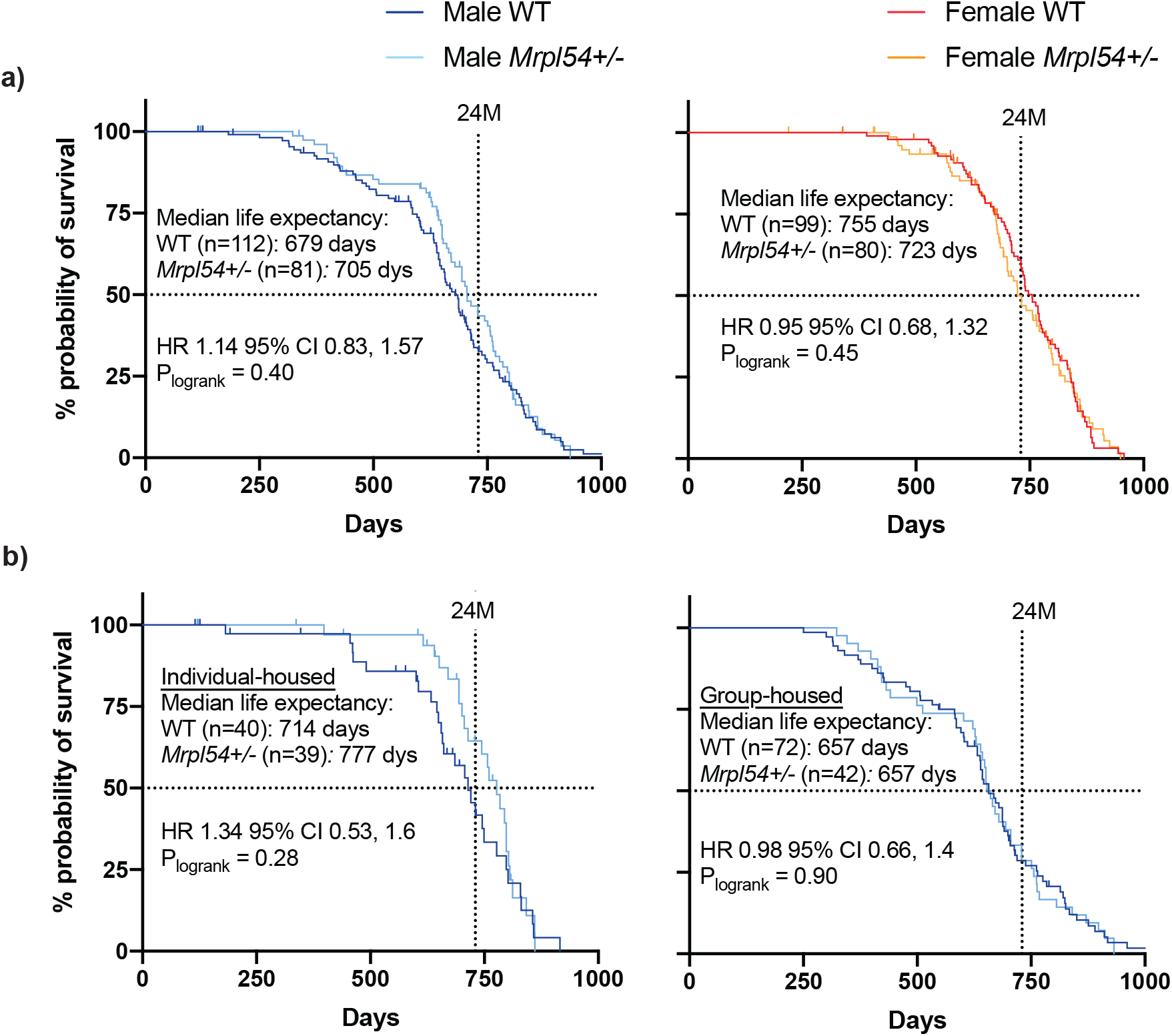
Life expectancy and lifespan in male and female *Mrpl54*^*+/-*^ mice. **a)** Kaplan Meier survival curves showed no difference in overall lifespan (top 10% of surviving animals) between either male *Mrpl54*^*+/-*^ (n=81) and WT (n=112) or female *Mrpl54*^*+/-*^ (n=80) and WT (n=99) mice. No differences were observed in median life expectancy between female *Mrpl54*^*+/-*^and WT; although, male *Mrpl54*^*+/-*^ showed a trend towards increased median life expectancy compared to male WT. **b)** Kaplan Meier survival curves showed no difference in overall lifespan between either individual-housed male *Mrpl54*^*+/-*^ (n=39) and WT (n=40) or group-housed male *Mrpl54*^*+/-*^ (n=42) and WT (n=72). There was a trend towards greater median life expectancy in individual-housed *Mrpl54*^*+/-*^ male mice compared to WT, with no difference in median life expectancy between group-housed male *Mrpl54*^*+/-*^ and WT.

During the lifespan study, some males were separated due to injuries or deaths from aggression and fighting, which resulted in a population of male mice that were housed individually (i.e., a single animal per cage from age 3-4 months onward). When individually and group-housed animals were analysed separately, it was evident that individually housed males had better median life expectancy than grouped-housed males for both genotypes, which is likely due to conspecific aggression increasing stress levels and shortening lifespan in group-housed males^42^. Amongst individually housed males, the *Mrpl54*^+/-^ animals exhibited a trend towards better median life expectancy (777 days vs. 714 days; Fig. 4b) and 24-month life expectancy compared to WT (24-month HR 1.85, 95% CI 0.86, 4.01, p = 0.12) (Fig. S4a). In contrast, group-housed males (n=2-9 males per cage) showed no differences in median or 24-month life expectancy between genotypes (Fig. S4b).

### Mrpl54^+/-^ did not activate the UPR^mt^ in liver mitochondria or primary myoblasts

Since MRP proteins play a crucial role in the translation of mtDNA-encoded mRNAs into protein subunits for oxidative phosphorylation (OXPHOS), we determined whether mitochondrial oxygen consumption was affected in the liver, a major metabolic organ. Using a Clark-type oxygen electrode, purified mitochondria from the liver of 7-week-old WT and *Mrpl54*^+/-^ males demonstrated similar rates of oxygen consumption (Fig. 5a). We then examined if reductions in *Mrpl54* induced a mitonuclear protein imbalance and induced the UPR^mt^. Using purified mitochondria from the liver of 7-week-old WT and *Mrpl54*^+/-^ males, we found no significant difference in HSP60 (HSPD1) protein levels, a classical marker for UPR^mt^ activation^43,44^ (Fig. 5b). This result was confirmed by mass spectrometry-based proteomics on purified liver mitochondria from 7-week-old males showing no differences in the levels of nDNA- or mtDNA-encoded proteins between WT and *Mrpl54*^+/-^ animals (Fig. 5c). Combined, these results suggest that the mitochondrial translational capacity was not affected in the liver tissue of *Mrpl54*^+/-^ animals.

**Figure 5.**
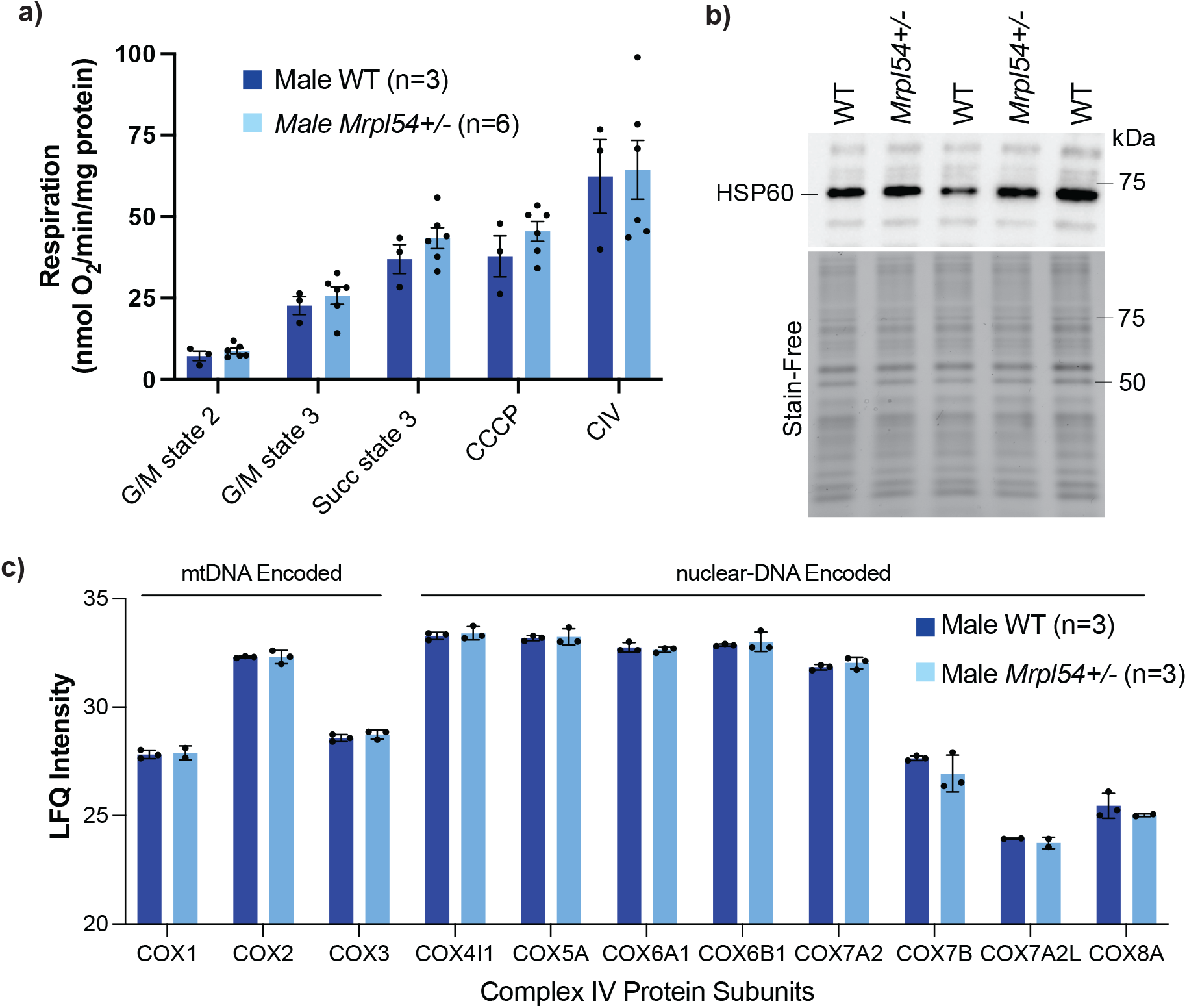
Biochemical profiling of Mrpl54+/- and wild type mouse liver. **a)** Oxygraph respiration analysis of isolated liver mitochondria showed no difference in state 2, state 3, or maximal respiration between *Mrpl54*^*+/-*^ (n=6) and WT (n=3). **b)** Semiquantitative western blot analysis of HSP60 expression in isolated liver mitochondria showed no difference between *Mrpl54*^*+/-*^ (n=2) and WT (n=3). **c)** Mass spectrometry analysis of protein from isolated liver mitochondria showed no difference between levels (label-free quantification, LFQ) of nDNA- and mtDNA-encoded protein subunits of ETC complex IV between *Mrpl54*^*+/-*^ (n=3) and WT (n=3). Graphs show mean ± SEM; male WT (blue); male *Mrpl54*^*+/-*^ (light blue)

Since the relative *Mrpl54* expression is highest in skeletal muscle tissue (Fig. 1c), we examined the effect of reduced *Mrpl54* expression on the levels of OXPHOS protein subunits and the UPR^mt^ marker HSP60 in primary myoblasts. Primary myoblast cultures were established from 7-week-old male *Mrpl54*^*+/-*^ and WT hindlimb skeletal muscle. In cell lysates from these myoblasts, levels of mitochondrial OXPHOS subunits (including both nDNA- and mtDNA-encoded proteins) were lower in *Mrpl54*^*+/-*^ mice compared to WT (Fig 6a). The level of succinate dehydrogenase complex iron sulphur subunit B (SDHB), a nDNA-encoded protein subunit of Complex II of the ETC, and the level of MTCO1, a mtDNA-encoded protein subunit of Complex IV of the ETC, were both lower in *Mrpl54*^*+/-*^ primary myoblasts (Fig 6a). Notably, considering that both MTCO1 and SDHB were similarly decreased, the ratio of both was not different in *Mrpl54*^*+/-*^ myoblasts. In line with this, HSP60 levels in *Mrpl54*^*+/-*^ myoblasts were not increased, suggesting that UPR^mt^ was not invoked in *Mrpl54*^*+/-*^ myoblasts (Fig 6b). These results suggest that reduced *Mrpl54*^*+/-*^ expression in primary myoblasts lowered levels of various ETC complexes but this reduction was insufficient to activate the UPR^mt^.

**Figure 6.**
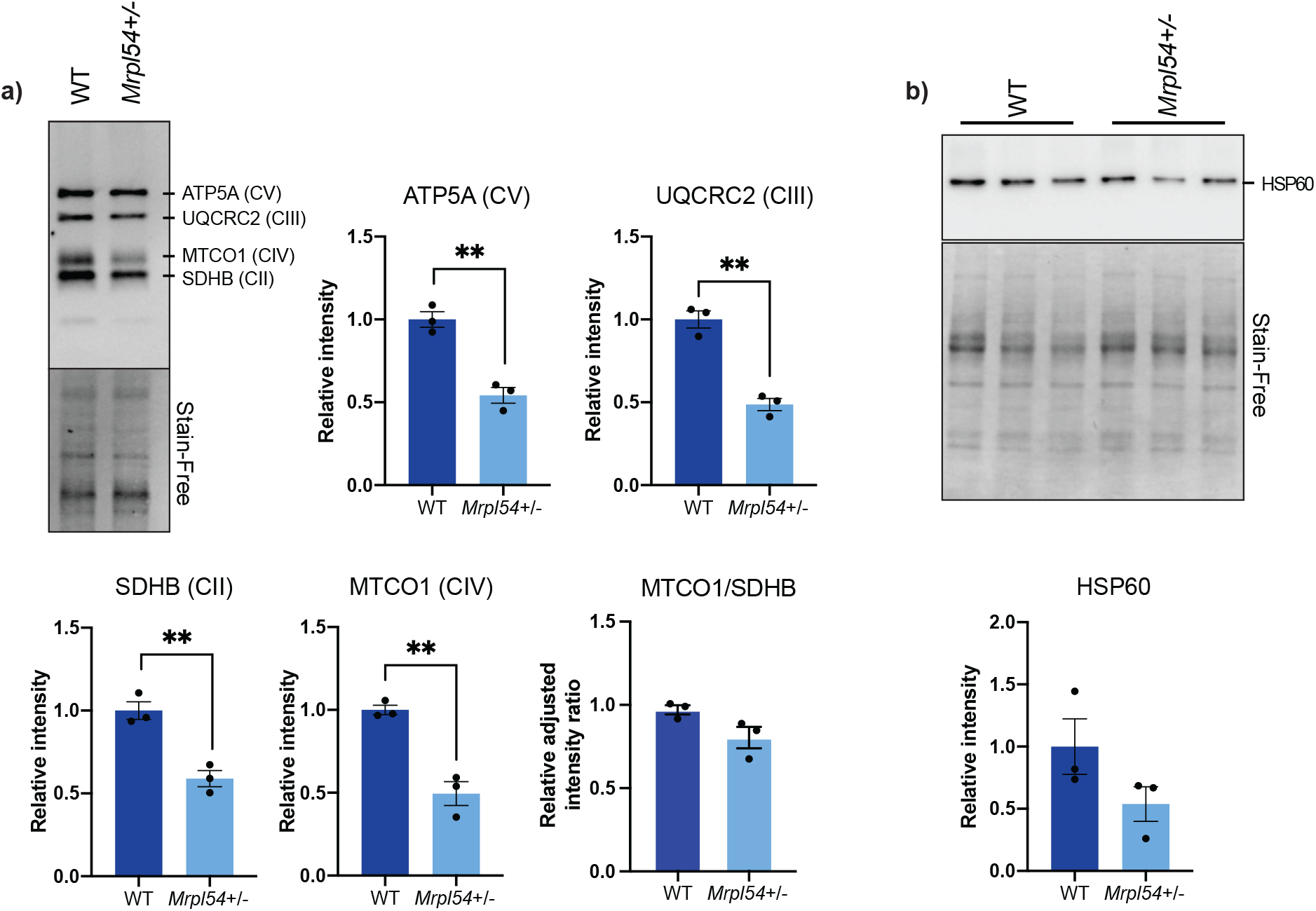
Biochemical profiling of *Mrpl54*^*+/-*^ and wild type mouse muscle. **a)** Semiquantitative representative western blot of ETC nDNA-encoded subunits (ATP5A, QCRC2, and SDHB) and mtDNA-encoded MTCO1 subunit in cell lysate from primary myoblasts showed a decrease in ETC subunit expression in *Mrpl54*^*+/-*^ (n=3) compared to WT (n=3). **b)** Semiquantitative western blot analysis of HSP60 expression in cell lysate from primary myoblasts showed no difference in HSP60 levels in *Mrpl54*^*+/-*^ (n=3) compared to WT (n=3). Graphs show mean ± SEM; male WT (blue); male *Mrpl54*^*+/-*^ (light blue); *P ≤ 0.05, **P ≤ 0.01, and ***P ≤ 0.001

## DISCUSSION

Reduced expression of MRP proteins in *C. elegans* results in an increase in both healthspan and lifespan through the induction of the UPR^mt^ that results from a mitonuclear protein imbalance^15^. Similarly, long lifespan is correlated with low expression of *Mrps5* and other MRP genes in a genetic reference population of mice and elevated UPR^mt^ markers are observed in cell cultures of the long-lived Snell dwarf mouse strain^15,29^. The UPR^mt^ is also induced in mice treated with doxycycline^45^, an antibiotic that inhibits mitochondrial translation^46^, which potentially reduces energy expenditure^45^. To determine whether reduced expression of *Mrpl54* in heterozygote mice alters metabolism or lifespan, we compared metabolic health at 6-, 18-, and 24-months as well as assessed lifespan and life expectancy in *Mrpl54*^+/-^ and WT animals.

While *Mrpl54* mRNA expression was reduced in multiple metabolic organs from *Mrpl54*^+/-^ animals, we found no compelling metabolic changes in 6-month-old adult mice. However, in line with previous reports showing an increase in rearing activity during test cage monitoring in mice treated with doxycycline^45^, female *Mrpl54*^+/-^ mice exhibited an increase in ambulatory activity along with elevated treadmill endurance when compared to WT. Like doxycycline-treated mice^15,45^, we observed a lower VO_2_ in adult female *Mrpl54*^*+/-*^ mice, but this is likely explained by their reduced body weight^40^, and not consistent with observations in male mice, or female mice at later time points.

We compared the life expectancy and lifespan in large cohorts of *Mrpl54*^+/-^ and WT male and female mice. As a result of injury or death associated with male aggression, the male cohort was divided into individual-housed and group-housed cages. Male mice tended to experience stress associated with conspecific aggression under group housing conditions. Male *Mrpl54*^*+/-*^ mice housed individually trended better in both median and 24-month life expectancy when compared to individual-housed male WT mice. In contrast, group-housed males had lower life expectancy compared to individual-housed males, with no differences between *Mrpl54*^+/-^ and WT. Male C57BL/6 mice can adapt to both individual as well as group housing conditions, the latter if they are housed together in low numbers^42^. As group-housed males tended to experience stress associated with conspecific aggression, it may be that the continuous stress experienced in group housing overcame any survival advantage afforded by the *Mrpl54*^+/-^ genotype.

Finally, we observe neither a mitonuclear imbalance nor an induction of the UPR^mt^ in *Mrpl54*^+/-^ mouse liver, a major metabolic organ. Within primary myoblasts, however, there was evidence of decreased OXPHOS protein expression, but no evidence of UPR^mt^ induction. It remains to be seen whether there is an induction of the UPR^mt^ in other *Mrpl54*^+/-^ mouse tissues and organs, or in a temporal fashion – neither of which was captured in this study. Previous work demonstrated that reductions in MRP expression (through RNAi of MRP genes, or inhibition of mitochondrial translation through doxycycline treatment) can increase healthspan and lifespan in *C. elegans* through a mitohormetic response^15^. Our work demonstrates that a reduction in *Mrpl54* expression through the germline may have implications in energy expenditure and activity levels in adult female mice, along with a potential increase in median and 24-month life expectancy of individual-housed male mice that are removed from the conspecific stress associated with group housing conditions. However, our findings also demonstrate that a reduction of *Mrpl54* is not sufficient to drive the induction of UPR^mt^ in mouse liver tissue or primary myoblasts, nor is it able to alter metabolism of older mice or affect lifespan.

It may be that reducing *Mrpl54* expression through the mammalian germline allows the mouse to adjust to lower MRPL54 protein levels through adaptations during developmental processes from embryogenesis onward – for example, genetic compensation where mutant mRNA degradation leads to transcriptional adaptation^47^. In other words, the levels of MRP expression are adjusted early in development to optimize mitochondrial output and maintain mitochondrial protein homeostasis. Alternatively, our negative results may indicate that the level of mitochondrial translation stress instigated by the *Mrpl54*^*+/-*^ genotype is not high enough to generate a mitonuclear imbalance, mito-cytosolic translational imbalance, or induce mitochondrial stress responses like the UPR^mt^ or ATF4-dependent signaling^15,21,22^. This explanation implies that any mitohormetic effect that affects metabolic health or lifespan requires a mitochondrial translational stress level above a specific threshold, and that this threshold was not met by the *Mrpl54*^*+/-*^ model.

In conclusion, our results demonstrate that the reduction of *Mrpl54* expression through the mouse germline does not induce a mitonuclear imbalance or a subsequent UPR^mt^ response. Furthermore, reduction in *Mrpl54* expression does not result in an improved life expectancy or overall lifespan. We recommend that future work pursue the hypothesis of a minimum threshold of mitochondrial translational stress necessary to invoke a mitohormetic impact on metabolic health with age or lifespan. As well, since reduced *Mrpl54* expression in male mice may mitigate the lifespan effects of stressful housing conditions, we recommend that the *Mrpl54*^*+/-*^ model be tested under various stress conditions, such as genotoxic or metabolic stress through a high fat diet.

## METHODS

### Animal Breeding

The Institute Clinique de la Souris (ICS) generated a *Mrpl54* constitutive knock-down mouse model on a C57BL/6N(Taconic) background (C57BL/6NTac-Mrpl54^tm1b(EUCOMM)Wtsi/IcsOrl^; colony name ICS-EPD1076_5_A09)^48,49^. *Mrpl54*^+/-^ breeding pairs were kept in ventilated racks and their progeny were housed in groups of 2-5 mice until their transfer to the metabolic health and longevity studies. Litter sizes were small (mean 4-5 pups) and production dropped off after the fourth litter. To generate the cohorts needed, CD1 host mothers were used for three rounds of *in vitro* fertilization, in addition to traditional breeding. At age 21-days, animals were weaned, ear-tagged, and tissue sampled from the ear for genotyping.

### Genotyping

DNA for genotyping was extracted from the ear tissue sample according to the Phire Animal Direct PCR protocol (F-170, Thermo Scientific). Genotyping proceeded according to the ICS protocol (Forward primer Ef 4877 5’-GACCCACATAAGCAGGGAAGGAGATG-3’, reverse primer L3r 4879 5’-CAATCTCCTGAGAATGTAGCCCACCAT-3’, Invitrogen). The *Mrpl54* knock-out allele generated a 402 base-pair (bp) fragment, and the WT allele generated a 1095 bp fragment.

### Animal husbandry

WT and *Mrpl54*^+/-^ animals, separated by sex, were housed under a 12:12-hour light-dark cycle in a dedicated room set to 23°C. All mice were fed a standard chow diet (Envigo; Teklad Global 2018; 18.6% crude protein, 6.2% fat, 44.2% available carbohydrates). Female and male mice were initially housed 5-10 per cage in a large rat cage (at least 100cm^2^ per mouse) to reduce stress^50^. Male cages experienced aggression and fighting and the male mice in cages that experienced injuries or deaths due to male-on-male aggression were reassigned to individual-housing. Animals in the metabolic health study were handled or weighed at least once per week. Animals in the longevity study were weighed once per month until age-related weight loss was observed, after which mice were weighed weekly or daily as needed.

### RNA isolation and Real-time qPCR

For total RNA isolation, a small piece of snap frozen mouse tissue was placed in 1ml TRIzol (Sigma-Aldrich or Invitrogen). Samples were homogenized with a 5mm steel bead using a TissueLyser II (Qiagen) for 3min at a frequency of 30Hz. RNA was isolated according to manufacturer’s protocol. From 1μg of extracted RNA, genomic DNA was eliminated with a gDNA Wipeout Buffer and subsequently reverse transcribed into cDNA using the QuantiTect Reverse Transcription Kit (Qiagen). The diluted cDNA was prepared with SYBR Green (Roche), and the quantitative gene expression was measured using the LightCycler 480 Instrument (Roche). The primer pairs used were as follows: *Mrpl54* forward 5’-AAAAAGCCAGTTGGCAAGGG-3’, reverse 5’-ATGTGTGGTGAGCTGAGTGG-3’; *36B4* (acidic ribosomal phosphoprotein gene) forward 5′-ACGGGTACAAACGAGTCCTG-3′, reverse 5′-GCCTTGACCTTTTCAGCAAG-3′; and *B2m* (beta2 microglobulin) forward 5′-GGCTCACACTGAATTCACCC-3, 5′-GTCTCGATCCCAGTAGACGG-3′. To normalize tissue sample expression (threshold cycle [Ct] values) to the expression of housekeeping genes, the ΔΔCt method was used. The geometric means of *36B4* and *B2m* genes were used to normalize *Mrpl54* tissue sample expression (ΔCt values) to the WT samples.

### Metabolic Health Analyses

The metabolic health of male and female WT and *Mrpl54*^+/-^ mice (n=7-17 in each group) were tested in three separate cohorts at 6-, 18-, and 24-months of age. The battery of tests started with EchoMRI body composition (EchoMRI-700 Analyzer), followed by indirect calorimetry using the comprehensive laboratory animal monitoring system (CLAMS, Columbus Instruments), oral glucose tolerance, intraperitoneal insulin tolerance, endurance treadmill, cold tolerance, heart rate and blood pressure measurement (at 18- and 24-months), and then necropsy (Fig. S1). Mice were allowed to recover 1-2 weeks between tests.

### EchoMRI and Indirect Calorimetry

Each mouse was loaded into an A100 antenna insert and placed into the EchoMRI (EchoMRI-700 Analyzer) to determine body composition (whole-body, lean, and fat mass). Immediately afterward, mice were placed in individual CLAMS (Columbus Instruments Oxymax System) plexiglass cages for 48-72 hours. The ambient temperature was set to thermoneutrality (28°C), and the light-dark cycle was 12:12. Mice had *ad libitum* access to water and powdered chow (Teklad 2018 Diet). At regular intervals (every 18-26 minutes), the following measurements were recorded: O_2_ volume (VO_2_), CO_2_ volume (VCO_2_), respiratory exchange ratio (RER), ambulatory and rearing motion, and the amount of food consumed in grams.

### Oral Glucose Tolerance Test (OGTT)

Mice were fasted overnight for 16-hours prior to the OGTT. Whole-blood glucose levels were measured using the Accu-check Performa glucometer and accompanying strips (Roche Diagnostics, Canada). For the 6M cohort, mice were placed in a restraint tube during the blood sampling from the left lateral tail vein. For the 18M and 24M cohorts, mice remained unrestrained on the home-cage hopper during blood sampling. At time 0, freshly made 20% α-D-glucose (Sigma-Aldrich PN158968) was orally administered using a flexible straight metal gavage at a dose of 1.0g per kg of body mass. Blood glucose was measured at baseline (prior to oral glucose dose), and at 15-, 30-, 45-, 60-, 90-, 120-, 150-, and 180-minutes post-gavage.

### Intraperitoneal Insulin Tolerance Test (iITT)

Two weeks prior to the iITT, intraperitoneal insulin titrations were conducted on a subset of mice to determine the correct dose to elicit an insulin response. Mice were fasted in the morning for 5-6 hours prior to the iITT. Whole-blood glucose levels were measured using the Accu-check Performa glucometer and accompanying strips (Roche Diagnostics, Canada). During blood sampling from the right lateral tail vein, mice remained unrestrained on their home-cage hopper. At time 0, mice received an intraperitoneal injection of insulin (NovoRapid Insulin aspart, 100U/mL, DIN 02245397). For the 6M cohort, females received an insulin dose of 1.5U/kg body mass and males received 2.5U/kg. For the 18M cohort, females received an insulin dose of 2U/kg and males received 4.5U/kg. For the 24M cohort, females received an insulin dose of 1U/kg and males received 2U/kg. Blood glucose was measured at baseline (prior to insulin injection), and at 15-, 30-, 45-, 60-, 75-, 90-, 105-, and 120-minutes post-injection. In the event the blood glucose level dropped below 2.0mmol/L, emergency α-D-glucose (Sigma-Aldrich PN 158968) was orally administered at a dose of 1g per kg of body mass.

### Treadmill

Mice were acclimated to the treadmill (Exer-3/6, Columbus Instruments) over three consecutive days where they were placed on an unmoving belt for 5 minutes, then ran at a constant speed of 15cm/s at an elevation of +5° with an electro-stimulus (0.1mA, 1Hz) applied to the resting grid. Those mice that achieved fewer than 5 electro-stimulations over 5 minutes by the third acclimation day were permitted to continue with the endurance test. The endurance test was conducted the day following the last day of acclimation using the following profile – accelerate to 15cm/s in 30 seconds and then run at 15cm/s for 12 minutes, accelerating +3.0cm/s over 30 seconds every 12 minutes^51^. Mice ran until the exhaustion criterion was met (5 electro-stimulus contacts [0.1mA, 1.0Hz] within 5 seconds) at which point the electro-stimulus was stopped, and the running time and maximum speed attained were recorded

### Cold Tolerance

The baseline rectal temperature of each mouse was measured using a lubricated rectal thermometer at an insertion depth ∼2cm. Mice were placed in individual cages with access to fresh water and a plastic house, but without access to bedding or food, and then placed in a +4°C refrigerated room with all cages at the same height. Rectal temperature was measured every hour for up to 4-6 hours. If rectal temperature reached 25°C or below^52^, the mouse was removed from the cold room and placed in a 37°C incubator with access to food and water.

### Heart Rate & Blood Pressure

Mice in the 18M and 24M cohorts were acclimated to the blood pressure and heart rate monitoring equipment (BP-2000-M-6 Series II Blood Pressure Analysis System, Visitech Systems) over a period of 3-5 days^53^. The procedure for acclimation and testing was the same – mice were individually placed in an opaque mouse holder on a warmed 36°C plate with the tail pulled through an inflatable tail cuff. Mice rested in position for 5 minutes, followed by 5 preliminary readings and then 10 actual readings. The program parameters were as follows: 2.5-second pause between readings, 10-second analysis pulse, system maximum set to 170mmHg. Test day immediately followed the last day of successful acclimation, where blood pressure was recorded for at least 6 out of 10 readings. The outcomes recorded were diastolic blood pressure, systolic blood pressure, and heart rate.

### EchoMRI & Necropsy

The day before the necropsy, body composition was determined using the EchoMRI-700 as described above. The mice were fasted overnight for 10-12 hours and then refed 1-2 hours before the necropsy. Mice were euthanized through injection with ketamine/xylazine cocktail (0.1mg/kg body weight) followed by exsanguination through cardiac puncture. The following tissues and organs were collected, weighed, fast-frozen in liquid N_2_ and then stored at -80°C for further analysis: pancreas, spleen, diaphragm, heart, liver, kidneys, quadriceps, gastrocnemius, soleus, tibialis anterior, and brain^54^. For the 24-month cohort necropsy, the right kidney, quadriceps, and gastrocnemius/soleus were suspended in Optimal Cutting Temperature fixative (Fisher Health Care, Cat# 4585) and placed on dry ice (kidney) or in isopentane (Sigma, CAT# 1003301470) supercooled in liquid N_2_ (muscles), then stored at -80°C for further analysis.

### Longevity Study and Growth

Male and female WT and *Mrpl54*^+/-^ mice (n=80-112 per group) were initially group-housed 5-10 per cage (at least 100cm^2^ per mouse). Male cages experienced aggression and fighting from age 3-months onward. In case of a fight-related injury or death, the males were reassigned to individual cages to improve animal welfare. Mice were weighed monthly, and median life expectancy (50% survival time point), 24-month life expectancy (all mice alive at 24 months were censored), and maximum lifespan (lifespan of top 10% of surviving animals) were calculated. Where a human-endpoint (HEP) was identified and time-permitting, mice were necropsied to identify tumour burden and to collect tissue and plasma for future analysis. HEPs included (a) >15% weight loss over 2 weeks, (b) skin lesions not responding to treatment, (c) head tilt/rolling or loss of balance, (d) ulcerated or bulging eye, (e) ulcerated mass, (f) respiratory distress, (g) inability to express bladder, or (h) a distended abdomen or growing internal mass^50^. Mice that were euthanized because of ulcerative dermatitis that did not respond to treatment (a C57BL/6 strain-specific condition^55^), or because of an unacceptable pain level from an ulcerated or bulging eye, were censored from the longevity analysis.

To assess growth over time, the body mass of WT and *Mrpl54*^+/-^ male and female animals were weighed monthly. Animals destined for the 6-, 18-, and 24-month metabolic health analyses were included in the growth analysis, up until the month they entered the metabolic health study.

### Respiration of Isolated Liver Mitochondria

7-week-old male WT and *Mrpl54*^+/-^ mice were fasted overnight for 12 hours, killed by cervical dislocation, and their livers weighed then rapidly dissected directly into ice-cold isolation buffer (300mM D-sucrose, 10mM Tris-HCl, 1mM EGTA-Tris Base; pH 7.2)^56^. The suspension was homogenized in a 15mL Potter-Elvehjem homogenizer, and the supernatant centrifuged twice at 1000*g* for 10 min at 4°C (Sorvall RC-6 Plus Centrifuge, Thermo Scientific). The supernatant was centrifuged 8000*g* for 10 min at 4°C and then aspirated without disturbing the pellet. The pellet was gently resuspended in ice-cold suspension buffer (300mM D-sucrose, 10mM Tris-HCl, 0.05mM EGTA-Tris Base; pH 7.2) and centrifuged at 8000*g* for 10 min at 4°C. The final pellet was gently re-suspended in 300μL suspension buffer and kept on ice.

The respiratory function of isolated liver mitochondria was measured using a Clark-type electrode (Oxygraph System, Hansatech Instruments Ltd.). The chambers were calibrated at 23°C with continuous stirring with a PTFE-coated magnetic follower bar. The mitochondrial suspension was quantified (Bio-Rad DC Protein Assay, BMG Labtech POLARstar Omega plate reader) and then suspended in miR05 respiration buffer (110mM D-sucrose, 60mM lactobionic acid, 20mM taurine, 20mM HEPES, 10mM KH2PO4, 3mM MgCl2, 0.5mM EGTA, 1g/L fatty-acid free BSA; pH 7.1 at 23°C)^57^ at 0.5mg protein/mL. Following a baseline recording, O_2_ consumption rate was measured following sequential additions of: 1) glutamate + malate (G/M, 5:2.5mM) to give Complex I (CI)-driven State 2 respiration, 2) ADP (2mM) to give CI-driven State 3 respiration, 3) Amytal (2mM) to inhibit CI, 4) succinate (10mM) to give Complex II (CII)-driven State 3 respiration, 5) carbonyl cyanide m-chlorophenylhydrazone (CCCP) titrations to decouple OXPHOS until maximum respiration was reached (0.01-0.05μM); 6) antimycin-A (8μM) to inhibit Complex III (CIII); 7) N,N,N’,N’-Tetramethyl-p-phenylenediamine dihydrochloride (TMPD) + ascorbate (5:0.3mM) to test Complex IV (CIV) respiration; and finally 8) KCN (0.6mM) to inhibit CIV.

### Mass Spectrometry (MS)-based Proteomics

Crude mitochondria were isolated from the livers of WT (n=3) and *Mrpl54*^+/-^ (n=3) male mice, as described above, and then purified using 30% Percoll gradient ultracentrifugation at 95,000*g* for 30 minutes at +4°C (Beckman sv41 rotor). Purified mitochondria were treated 1:200 with a protease inhibitor cocktail (Thermo Fisher Scientific HALT PI) then the mitochondrial proteins were left to precipitate overnight in precipitation buffer (50% acetone, 49.9% ethanol, 0.1% acetic acid). Proteins were suspended in 50mM ammonium bicarbonate in 8M urea and then quantified (Bio-Rad DC Protein Assay, BMG Labtech POLARstar Omega plate reader). To reduce disulphide bonds, 140μg of mitochondrial protein was incubated with 1:100 of 1M dithiothreitol (Sigma-Aldrich, SKU 43815), followed by 1:50 dark incubation in 1M iodoacetamide (Sigma-Aldrich, SKU I1149). The urea concentration was reduced to <2M and the protein sample was digested overnight at 37°C with 1:100 Trypsin Gold (Promega V5280). The following day, the digested proteins were passed through a Sep-Pak Vac 1cc tC18 cartridge (Waters, WAT054960) to rid the sample of salts and contaminants. The sample was dried in a Savant DNA120 SpeedVac Concentrator (Thermo Electron Corporation) at room temperature and analysed using a nano-LC-MS/MS with an Orbitrap Elite mass spectrometer (Thermo Fisher Scientific) coupled to an ultraperformance LC system (Ultimate 3000 RSLC; Thermo Fisher Scientific). Data analysis was performed with Proteome Discoverer (version 1.3), and searches were performed with Mascot and Sequest against a mouse database (UniProt). Data were further processed, inspected, and visualized with Scaffold 4 (Proteome Software).

### Generation of Primary Myoblasts

Hind limb skeletal muscles were excised from euthanized 7-week-old male WT (n=3) and *Mrpl54*^+/-^ (n=3) mice. Under aseptic conditions, the muscles were rinsed in phosphate buffered solution (PBS, Wisent Inc., Cat# 311-010-CL), the fat excised, and the remaining muscle tissue placed in collagenase B-dispase solution, 1.5mL per 0.5g tissue (4mg/mL dispase II [Sigma D4693], 10mg/mL collagenase B [Roche 11088831001] in Hams-F10 [Multicell, Wisent Inc. Cat# 318 050-CL]). The tissue was mulched with a razor blade, incubated at 37°C for 30 minutes, and then homogenized through trituration until smooth. The homogenate was passed through a 100micron cell strainer with PBS washes. The muscle cells were centrifuged at 300*g* for 5 minutes and the cell pellet resuspended in Ham’s Complete growth medium (4% bovine calf serum [cytiva, Cat# SH30077.03], 1% penicillin and streptomycin [gibco by Life Technologies, Cat# 15140122], 2.5ng/μL human basic fibroblast growth factor [bFGF, Stemcell, Cat# 78003]). To rid the culture of fibroblasts, the cells were allowed to rest on a non-collagen-coated plate for two hours at 37°C before plating onto a 10cm collagen-coated plate. To enrich myoblasts, 80% confluent plates were pre-plated on non-collagen-coated plates for one hour before plating on a collagen-coated plate.

### Western Blotting

Protein was extracted by suspending isolated WT and *Mrpl54*^+/-^ liver mitochondria or primary myoblast cells (10cm plates at 80% confluence) in radioimmunoprecipitation (RIPA) lysis buffer (50mM Tris HCl, 150mM NaCl, 1% v/v Triton X-100, 0.5% w/v sodium deoxycholate, 0.1% w/v SDS; pH 8.0) augmented with protease and phosphatase inhibitors (1:100 HALT PI cocktail, Thermofisher Sci Cat# 78440; or Roche cOmplete ULTRA [Cat# 05892970001] plus phosStop [Cat# 4906845001] tablets), then flash frozen in liquid N_2_. The solution was centrifuged 16,000*g* for 10 minutes at 4°C and the supernatant containing the mitochondrial or cell lysate proteins quantified (Bio-Rad DC Protein Assay, BMG Labtech POLARstar Omega plate reader).

Protein samples were diluted to 1μg/μL in Laemmli Buffer (Bio-Rad Cat# 1610737) augmented with 10% *β*-mercaptoethanol (Fisher Bioreagents, BP176-100) and run on a 10% SDS-PAGE using the TGX Stain-free FastCast Acrylamide Kit (Bio-Rad Cat# 1610183) and the Mini-PROTEAN system (Bio-Rad Cat# 1658006). SDS-PAGE was run at a constant 90 volts across the gel for 30 minutes and then 120 volts for 50-60 minutes in a 4°C cold room. The StainFree gels were activated once for 45 seconds using a ChemiDoc Touch Imaging System (Bio-Rad) and then the proteins transferred to TransBlot Turbo Mini size 0.2μm nitrocellulose membrane (Bio-Rad) using the Trans-Blot TurboTransfer System (Bio-Rad Cat# 1704150EDU).

The nitrocellulose membrane was imaged for total protein and then blocked for one hour in blocking buffer (5% w/v bovine serum albumin [Sigma, SKU A7906] in TBS-T buffer [50mM Tris-HCl, pH 7.6; 150mM NaCl; 0.1% Tween]), rocking gently at room temperature. Membranes were then incubated with primary antibody diluted in TBS-T buffer (1:1000 Total OXPHOS Rodent WB Antibody Cocktail [abcam Cat# ab110413] or 1:1000 HSP60 XP Rabbit mAB [Cell Signaling Technology CAT#12165]) overnight at 4°C with gentle rocking. After three washes with TBS-T, the membrane was incubated for one hour with the matching IgG, horseradish peroxidase (HRP)-conjugated secondary antibody (1:10,000 Cell Signaling Technology Anti-rodent Cat# 7076, or Anti-rabbit Cat# 7074P2), rocking at room temperature. After three washes in TBS-T, the membrane was visualized using enhanced chemiluminescent detection (BioRad Clarity Cat# 1705061) on the ChemiDoc Touch Imaging System (Bio-Rad).

### Statistical Analysis

All data are presented as mean ± SEM, unless otherwise stated. Differences between two groups were assessed using two-tailed t-tests. Analysis of covariance (ANCOVA) was used to eliminate unwanted variance on the dependent variable. To compare the interaction between time and treatment, a two-way analysis of variance (ANOVA) tests was performed. Survival curves were analysed using the Kaplan-Meier method. Hazard ratios were calculated using the Log rank test. GraphPad Prism 9.4.1 (GraphPad Software, Inc.) was used for all statistical analyses, and *p*<0.05 was considered significant.

## ACKNOWLEDGEMENTS

We thank the Institut Clinique de la Souris – PHENOMIN for the establishment of the *Mrpl54* mouse mutant line. From the University of Ottawa, we thank the following people: Adriana Gambarotta for her IVF expertise; Dr. Kerstin Ure, Mirela Barclay, and Sarah Kealey from the Behavioural Core Unit for help with CLAMS; the staff at the Animal Care and Veterinary Services Unit, in particular Courtney Reeks and Christine Kitchen, for help in mouse-handling training and colony maintenance assistance; and Prof. Daniel Figeys for mass spectrophotometry proteomics collaboration (NorthOmics). This project is financially supported by a VIDI grant (917.15.305) to RHH, UvA Lustrum 385 grant to EGD, NSERC CGS-D scholarship awarded to KR, and grants awarded to KJM.

## Author Contributions

Conception of hypotheses: R.H. Houtkooper. Design of the work: R.H. Houtkooper, K.J. Menzies, K. Reid, and E.G. Daniels. Acquisition and analysis of the data: K. Reid, E.G. Daniels, R. Kamble, G. E. Janssens, M. Hu, and G. Vasam. Drafting of the manuscript: K. Reid, E.G. Daniels, A. Green, R.H. Houtkooper and K. Menzies. Analysis and interpretation of the data: K. Reid, K. Menzies, E.G. Daniels, G. Vasam, A. Green, and R.H. Houtkooper. All authors approved the final version of the manuscript.

