## Supplemental Figures S1-S4 for "Reducing mitochondrial ribosomal gene expression does not alter metabolic health or lifespan in mice"


Kim Reid* [1]

Eileen G. Daniels* [2,3]

Goutham Vasam[4]

Rashmi Kamble[2]

Georges E. Janssens[2,3]

Man Hu[2,3]

Alexander E. Green[4,5]

Riekelt H. Houtkooper** [2,3,6]

Keir J. Menzies** [4,5]

1. Department of Biology. University of Ottawa, 30 Marie Curie Private, Ottawa, Ontario, Canada
2. Laboratory Genetic Metabolic Diseases, Amsterdam UMC location University of Amsterdam, Meibergdreef 9, 1105 AZ Amsterdam, The Netherlands
3. Amsterdam Gastroenterology Endocrinology and Metabolism institute, Amsterdam, The Netherlands
4. Interdisciplinary School of Health Sciences, University of Ottawa, 451 Smyth Road, K1H 8M5 Ottawa, Ontario, Canada
5. Ottawa Institute of Systems Biology and the Éric Poulin Centre for Neuromuscular Disease, Department of Biochemistry, Microbiology and Immunology, Faculty of Medicine, University of Ottawa, Ottawa, Ontario, Canada
6. Amsterdam Cardiovascular Sciences institute, Amsterdam, The Netherlands

*co-first authors

**co-last authors

**Figure S1 | Protocol and timeline for metabolic phenotyping of *Mrpl54*$^{+/-}$ and WT mouse cohorts**

Metabolic phenotyping in 6-, 18-, and 24-month male and female cohorts began with an assessment of body composition (EchoMRI) followed by 24-48-hour measurement of $VO_2$, RER, and ambulatory motion (CLAMS). Each cohort was then assessed using the following tests with 1-2 weeks between each test, in order: blood glucose tolerance (OGTT), intraperitoneal insulin tolerance (iITT), cold challenge (4°C), heart rate and blood pressure monitoring (18M and 24M only), and endurance treadmill, ending with a necropsy and tissue, organ, and plasma collection. Images were created with BioRender.com.

**Figure S2 | Change in *Mrpl54*$^{+/-}$ and WT body composition at age 6-, 18-, and 24-months**

Body composition (g) by EchoMRI with increasing age for both male and female *Mrpl54*$^{+/-}$ and WT mice. Graphs show mean ± SEM; male WT (blue); male *Mrpl54*$^{+/-}$ (light blue); female WT (orange); and female *Mrpl54*$^{+/-}$ (light orange). $^{**}P \leq 0.01$, $^{***}P \leq 0.001$, and $^{****}P \leq 0.0001$

**Figure S3 | Body composition and metabolic testing results in 18M aged male and female mice: *Mrpl54*$^{+/-}$ versus WT**

**a)** Body composition (g) by EchoMRI in 18M aged mice showed no difference in total body, fat, or lean mass between either male *Mrpl54*$^{+/-}$ (n=11) and WT (n=13) or female *Mrpl54*$^{+/-}$ (n=12) and WT (n=12).

**b)** Mean $VO_2$ (L/hour) by CLAMS and body mass ANCOVA analysis (insets) in 18M aged mice showed no differences for 24-hour, light, or dark periods between either male *Mrpl54*$^{+/-}$ (n=11) and WT (n=12) or female *Mrpl54*$^{+/-}$ (n=12) and WT (n=12).

**c)** Mean RER by CLAMS in 18M aged mice showed no difference between either male *Mrpl54*$^{+/-}$ (n=11) and WT (n=12) or female *Mrpl54*$^{+/-}$ (n=12) and WT (n=12).

**d)** Mean ambulatory motion (beam break counts) in 18M aged mice showed no difference between either male *Mrpl54*$^{+/-}$ (n=11) and WT (n=12) or female *Mrpl54*$^{+/-}$ (n=12) and WT (n=12).

**e)** Mean duration (minutes) on an endurance treadmill for 18M aged mice showed no difference in endurance time between either male *Mrpl54*$^{+/-}$ (n=9) and WT (n=11) or female *Mrpl54*$^{+/-}$ (n=10) and WT (n=9).

**f)** Mean blood glucose level (mmol/L) over time in response to an oral glucose challenge (OGTT) in 18M aged mice showed no difference between either male *Mrpl54*$^{+/-}$ (n=11) and WT (n=12) or female *Mrpl54*$^{+/-}$ (n=11) and WT (n=12).

**g)** Mean blood glucose level (mmol/L) and AUC (insets) over time in response to intraperitoneal insulin challenge (iITT) in 18M aged mice showed no differences between either male *Mrpl54*$^{+/-}$ (n=12) and WT (n=13) or female *Mrpl54*$^{+/-}$ (n=10) and WT (n=11).

**h)** Mean rectal temperature (°C) in response to a 4-hour 4°C cold challenge in 18M aged mice showed no difference between either male *Mrpl54*$^{+/-}$ (n=10) and WT (n=9) or female *Mrpl54*$^{+/-}$ (n=10) and WT (n=9).

**i)** Mean heart rate (beats/min) and systolic blood pressure (mmHg) in 18M aged mice showed no differences between either male *Mrpl54*$^{+/-}$ (n=10) and WT (n=11) or female *Mrpl54*$^{+/-}$ (n=10) and WT (n=7). Diastolic blood pressure (mmHg) at 18M was higher in male *Mrpl54*$^{+/-}$ compared to WT, with no differences between female *Mrpl54*$^{+/-}$ and WT.

**j)** Relative organ weights (% body mass) in 18M aged necropsied mice showed no differences between either male *Mrpl54*$^{+/-}$ (n=7) and WT (n=12) or female *Mrpl54*$^{+/-}$ (n=10) and WT (n=7).

Graphs show mean ± SEM; *P ≤ 0.05, male WT (blue); male *Mrpl54*$^{+/-}$ (light blue); female WT (orange); and female *Mrpl54*$^{+/-}$ (light orange).

**Figure S4 | 24-month life expectancy in *Mrpl54*$^{+/-}$ versus WT males by housing conditions**

**a)** Kaplan Meier survival curves show no difference in 24-month life expectancy (where all mice alive after 24-month are censored) in group-housed males between *Mrpl54*$^{+/-}$ (n=42) and WT (n=72).

**b)** Kaplan Meier survival curves show a trend towards increased 24-month life expectancy in individual-housed *Mrpl54*$^{+/-}$ (n=39) males compared to WT (n=40).

Fig. S1

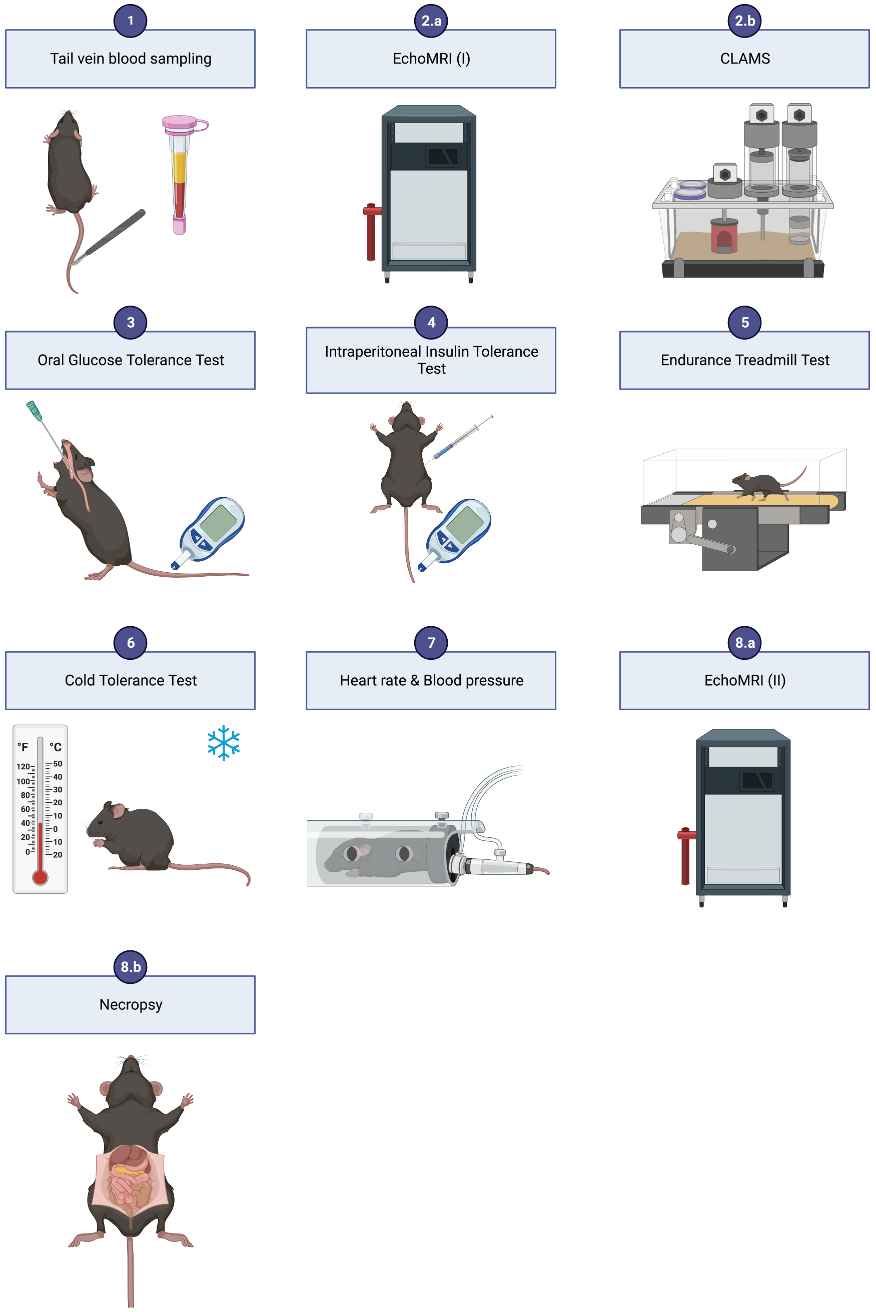
1
Tail vein blood sampling
2.a
EchoMRI (I)
2.b
CLAMS
3
Oral Glucose Tolerance Test
4
Intraperitoneal Insulin Tolerance Test
5
Endurance Treadmill Test
6
Cold Tolerance Test
7
Heart rate & Blood pressure
8.a
EchoMRI (II)
8.b
Necropsy

Fig. S2

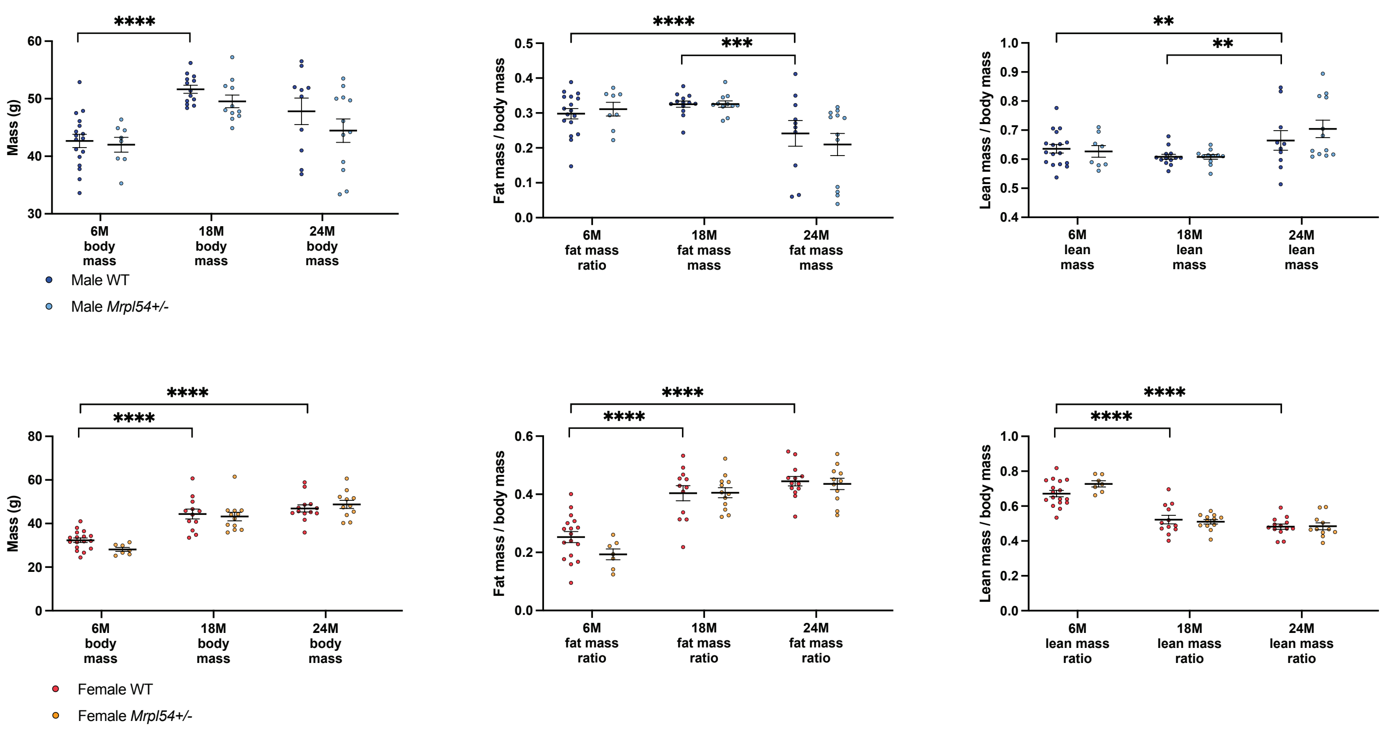
****
Mass (g)
60
50
40
30
6M body mass
18M body mass
24M body mass
Male WT
Male Mrpl54+/-
****
***
Fat mass / body mass
0.5
0.4
0.3
0.2
0.1
0.0
6M fat mass ratio
18M fat mass mass
24M fat mass mass
**
**
Lean mass / body mass
1.0
0.9
0.8
0.7
0.6
0.5
0.4
6M lean mass
18M lean mass
24M lean mass
****
****
Mass (g)
80
60
40
20
0
6M body mass
18M body mass
24M body mass
Female WT
Female Mrpl54+/-
****
****
Fat mass / body mass
0.6
0.4
0.2
0.0
6M fat mass ratio
18M fat mass ratio
24M fat mass ratio
****
****
Lean mass / body mass
1.0
0.8
0.6
0.4
0.2
0.0
6M lean mass ratio
18M lean mass ratio
24M lean mass ratio

Fig. S3

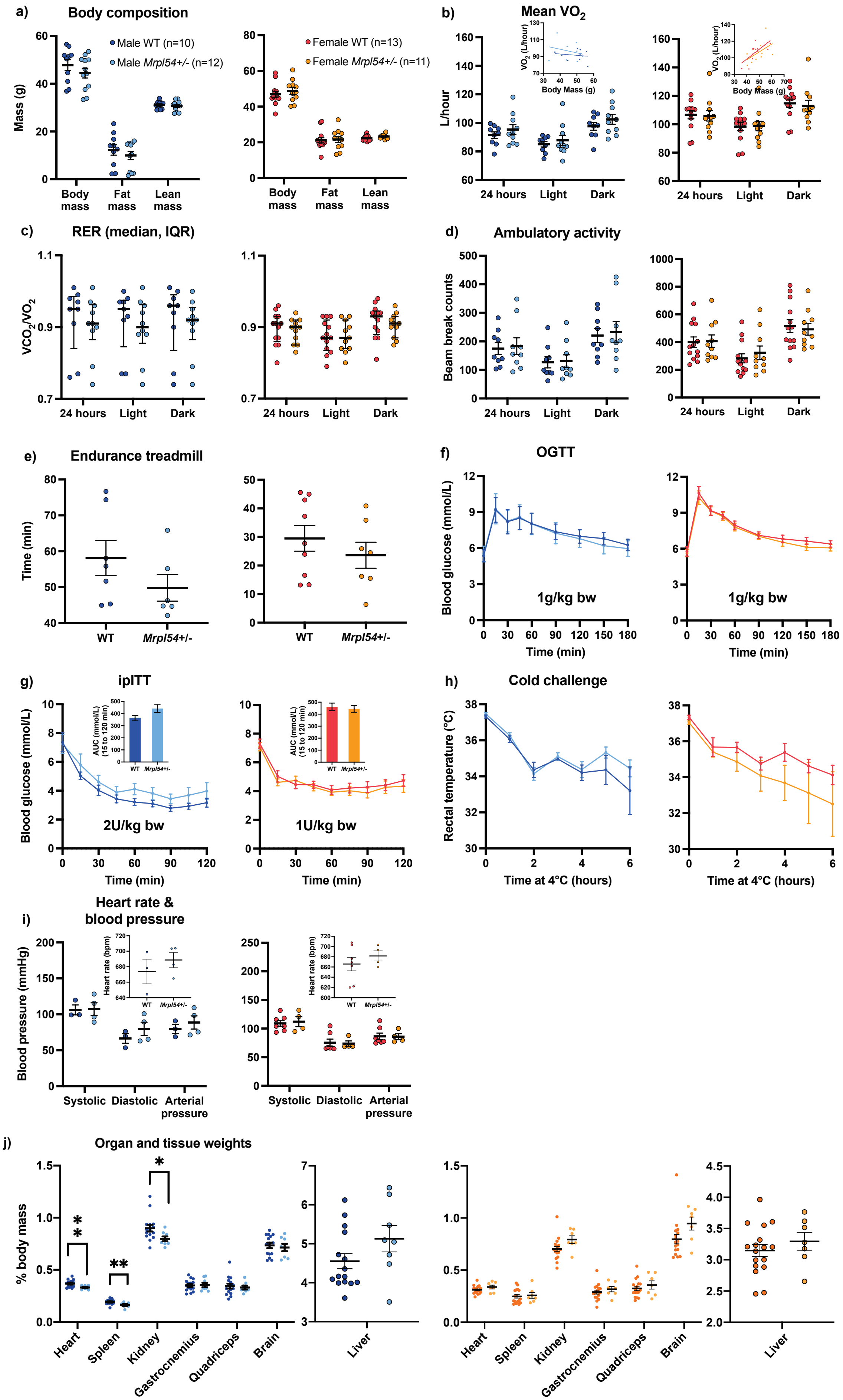

a) Body composition
Male WT (n=10)
Male Mrpl54+/- (n=12)
Female WT (n=13)
Female Mrpl54+/- (n=11)
Mass (g)
Body mass
Fat mass
Lean mass
b) Mean VO2
L/hour
VO2 (L/hour)
Body Mass (g)
24 hours
Light
Dark
c) RER (median, IQR)
VCO2/VO2
d) Ambulatory activity
Beam break counts
e) Endurance treadmill
Time (min)
WT
Mrpl54+/-
f) OGTT
Blood glucose (mmol/L)
1g/kg bw
Time (min)
g) ipITT
AUC (mmol/L) (15 to 120 min)
2U/kg bw
1U/kg bw
h) Cold challenge
Rectal temperature (°C)
Time at 4°C (hours)
i) Heart rate & blood pressure
Blood pressure (mmHg)
Heart rate (bpm)
Systolic
Diastolic
Arterial pressure
j) Organ and tissue weights
% body mass
Heart
Spleen
Kidney
Gastrocnemius
Quadriceps
Brain
Liver

Fig. S4

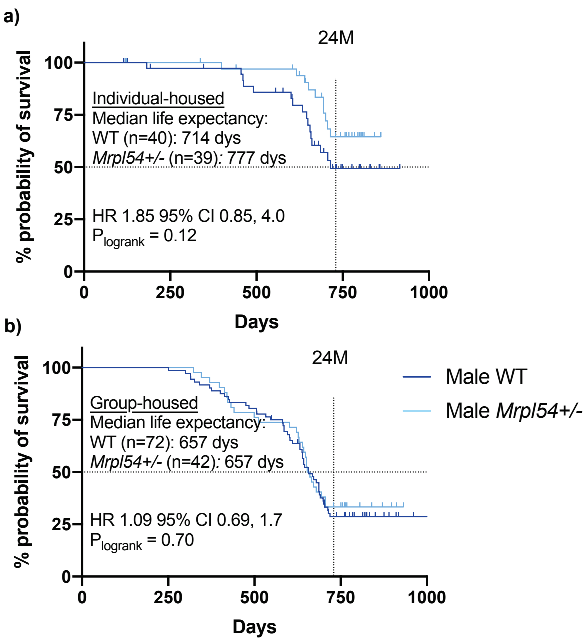

a)
24M
Individual-housed
Median life expectancy:
WT (n=40): 714 dys
Mrpl54+/- (n=39): 777 dys
HR 1.85 95% CI 0.85, 4.0
P logrank = 0.12
% probability of survival
100
75
50
25
0
0
250
500
750
1000
Days
b)
24M
Group-housed
Median life expectancy:
WT (n=72): 657 dys
Mrpl54+/- (n=42): 657 dys
HR 1.09 95% CI 0.69, 1.7
P logrank = 0.70
% probability of survival
100
75
50
25
0
0
250
500
750
1000
Days
Male WT
Male Mrpl54+/-
